# Investigating epithelial-mesenchymal heterogeneity of tumors and circulating tumor cells with transcriptomic analysis and biophysical modeling

**DOI:** 10.1101/2020.10.30.362426

**Authors:** Federico Bocci, Susmita Mandal, Tanishq Tejaswi, Mohit Kumar Jolly

## Abstract

**Introduction:** Cellular heterogeneity along the Epithelial-Mesenchymal Plasticity (EMP) spectrum is a paramount feature observed in tumors and circulating tumor cells (CTCs). High-throughput techniques now offer unprecedented details on this variability at a single-cell resolution. Yet, there is no current consensus about how EMP in tumors propagates to that in CTCs. To investigate the relationship between EMP associated heterogeneity of tumors and that of CTCs, we integrated transcriptomic analysis and biophysical modeling.

**Methods:** We apply three EMT (Epithelial-Mesenchymal Transition) scoring metrics to multiple tumor samples and CTC datasets from several cancer types. Moreover, we develop a biophysical model that couples EMT associated phenotypic switching in a primary tumor with cell migration. Finally, we integrate EMT transcriptomic analysis and *in silico* modeling to evaluate the predictive power of several measurements of tumor aggressiveness, including tumor EMT score, CTC EMT score, fraction of CTC clusters found in circulation, and CTC cluster size distribution.

**Results:** Analysis of high-throughput datasets reveals a pronounced heterogeneity without a well-defined relation between EMT traits in tumors and CTCs. Moreover, mathematical modeling predicts different phases where CTCs can be less, equally, or more mesenchymal than primary tumor depending on the dynamics of phenotypic transition and cell migration. Consistently, various datasets of CTC cluster size distribution from different cancer types are fitted onto different regimes of the model. By further constraining the model with experimental measurements of tumor EMT score, CTC EMT score, and fraction of CTC cluster in bloodstream, we show that none of these assays alone can provide sufficient information to predict the other variables.

**Conclusions:** By integrating analysis of single cell gene expression and *in silico* modeling, we propose that the relationship between EMT progression in tumors and CTCs can be variable, and in general, predicting one from the other may not be as straightforward as tacitly assumed.

## Introduction

Epithelial-Mesenchymal Transition (EMT) (now increasingly referred to as EMP - Epithelial-Mesenchymal Plasticity) is a crucial axis of tumor progression that regulates motility, resistance to therapies and proliferation [1]. During EMT, epithelial cancer cells can partially or completely lose their E-cadherin mediated cell-cell adhesion and apicobasal polarity, instead becoming motile, mesenchymal cells [2]. Recent experimental and computational findings suggest that EMT is not necessarily a binary process; instead cells can manifest one or more hybrid E/M phenotype(s) with mixed traits of epithelial and mesenchymal cells along the EMT (or EMP) spectrum [3–8].

Indeed, cellular heterogeneity is emerging as a hallmark of cancer, with cells with distinct EMT phenotypes often localized in distinct tumor regions [9,10]. Similarly, circulating tumor cells (CTCs) that launch into circulation to give rise to metastasis can exhibit a spectrum of EMT features [11]. Hybrid E/M cancer cells that partially conserve cell adhesion can travel the bloodstream collectively as highly metastatic CTC clusters [12]. Recent advances in high-throughput techniques such as RNA-sequencing are providing insights into the multi-faceted dynamics of EMT, predicting the number of intermediate hybrid E/M states and the most probable EMT trajectories [7]. However, this heterogeneity sparks questions about the relationship between EMT progression in primary tumors and CTCs. Is it possible to reliably predict the composition of a tumor from gene expression measurements of CTCs, and/or *vice versa*, or does the relationship between EMT progression in tumors and CTCs depend on tumor type, subtype, or even patient-specific factors? Furthermore, how much can be said about EMT progression in tumors by measuring statistical properties of CTC migration, such as the fraction of CTC clusters or CTC cluster size distribution from blood samples?

Here, we tackle these poignant questions by integrating transcriptomic analysis and computational modeling. First, we apply three EMT scoring metrics to several tumor and CTC datasets; these scores, which correlate well with one another, demonstrate that the EMT traits of tumors and CTCs are highly heterogeneous, raising questions about how much can be predicted about the EMT score of CTCs from the primary tumor and *vice versa*. To further investigate this heterogeneity and interdependence of EMT in tumors and CTCs, we turn to *in silico* biophysical modeling that couples EMT in the primary tumor and cell migration. The model reveals different parameter regions in which CTCs can either be more mesenchymal or more epithelial than the primary tumor, depending on the rate of EMT and migration dynamics (collective vs. individual). Moreover, several CTC cluster size distribution datasets sampled from different tumors are mapped onto different parametric combinations in the model description, suggesting that the heterogeneous tumor-CTC EMT relation could be an important aspect *in vivo*. Finally, we integrate the EMT scoring and biophysical model in a single computational pipeline to investigate how much can be predicted by this biophysical model, in terms of tumor and CTC EMT score, CTC cluster fraction and CTC cluster size distribution, when only one of these variables is provided as an input.

## Materials and methods

### 1. Data analysis

We calculated EMT scores for multiple primary tumor and CTC datasets using three different EMT metrics – 76GS, KS, and MLR [13–15]. We also calculated correlations in the EMT scores of samples for a given dataset using Spearman’s and Pearson’s correlation coefficient values.

### 2. EMT model

The biophysical model focuses on cells at the periphery of a tumor that have the potential to undergo EMT and migrate as circulating tumor cells (CTCs) individually or as a cluster. Therefore, cells in the model are arranged on a one-dimensional lattice that represents the tumor invading edge. Conversely, cells in the more internal layers are not modeled explicitly since they lack the physical space to migrate. Starting from an epithelial state, cells at the invading edge undergo transitions with a rate (k) through a number of intermediate hybrid E/M states, and eventually to a mesenchymal state.

Mesenchymal cells can migrate from the cell layer as individual cells; conversely, clusters of neighboring hybrid E/M cells can migrate together as multicellular units. The migration rate of clusters depends on i) the number of cells in the cluster (i.e. the cluster size), ii) the EMT state of neighboring cells as cell-cell adhesion bonds must be broken, and iii) a migration cooperativity parameter (c) that quantify the propensity to collective migration. While a low c favorites individual migration, a large c promotes clustered migration if hybrid E/M cells are in contact. The migration is simply modeled as a discrete event and the physical motion of the migrating cells is not considered explicitly. When a single cell or cluster migrates, cell(s) are instantaneously replaced by new epithelial cells. This process, which ensures a constant number of cells in the invading layer, considers the emergence of interior cells that become exposed to EMT-inducing signals once peripheral cells migrate.

The dynamics of cell fractions with a generic number (N) of hybrid E/M states are described by a set of ordinary differential equations:

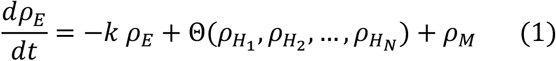

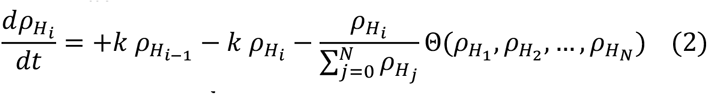

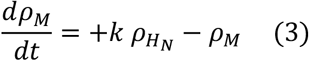

In Eqs. (1-3), k is an EMT rate in arbitrary, dimensionless units. Eq. 2 represents the dynamics of the i-th hybrid E/M state cell population, which is increased due to transitions from the i-1 state 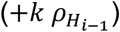 and decreased by transitions to the i+1 state 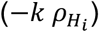. The supplementary section (**SI section 1.1**) shows the form of the equations for the case of three intermediate states (E-like, E/M, M-like) that is used throughout most of the paper. Furthermore, the term 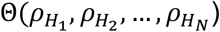 quantifies the loss of hybrid E/M cells due to migration out of the lattice (either as single cells or multicellular clusters); the explicit form of this term is derived in the supplementary information (**SI section 1.2-1.5**). The mesenchymal cell fraction (eq. 3) increases due to transition from the terminal hybrid E/M state 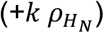 and decreases due to single cell migration (−*ρ*_*M*_). Since migrating cells are replaced by new epithelial cells, the epithelial cell fraction (eq. 1) increased via an influx equal to the migration loss 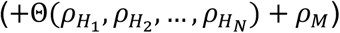. It is worth noting that all migration rates in Eqs. (1-3) are not explicitly multiplied by a migration rate constant; since the model is dimensionless, time is rescaled so that this parameter is equal to 1. Therefore, the EMT rate (k) can be thought of as a ratio between the EMT rate and cell migration rate.

Custom-made python scripts to solve the model and reproduce the results are freely available at https://github.com/federicobocci/Biophysical-model-of-EMT-heterogeneity.

## Results

### 1. EMT scoring metrics analysis reveals heterogeneity in primary tumors and CTCs across cancer types

Recent approaches have investigated the varying degrees of EMT in CTCs using a handful of markers and their association with patient survival across cancer types [16–20]. Further, there has been a surge of high-throughput measurement such as RNA-seq of CTCs [21–23] as well as primary tumor [24–26], including investigations at a single-cell level. Given the heterogeneity of assessing EMT in multiple studies using diverse markers [27], such transcriptomics-based measurements can enable quantifying EMT in a more systematic manner using different scoring methods.

We calculated the EMT scores of multiple publicly available datasets associated with CTCs, using three different EMT metrics – 76GS, KS and MLR [28]. These three methods use different gene lists and algorithms, and have been developed based on pan-cancer signatures of EMT identified from preclinical (*in vitro*) and/or clinical data [13–15]. Thus, these scores can indicate the extent of EMT a cell line, primary/metastatic tumor or CTC has undergone.

The more epithelial a sample is, the lower its KS score (on a scale of [-1, 1]) or MLR score (on a scale of [0,2]) and the higher is its 76GS score (no *a priori* defined range of values). Thus, for a given dataset, while we expected a positive correlation between MLR and KS scores, we expected 76GS scores to correlate negatively with KS and MLR ones.

EMT scores of 4 breast cancer CTC cell lines and their metastatic variants [29] (GSE112855) using the above-mentioned three metrics, displayed heterogeneity in their EMT-ness (**Fig 1A, S1A-E, S2A-C**). However, none of these cell/sub lines could be classified as strongly mesenchymal, based on scores across the three metrics. Next, CTCs from various breast cancer patients exhibited heterogeneity [30] (PRJNA471754); however, the samples were overall shifted towards a more mesenchymal end of the spectrum as compared to breast cancer cell lines (GSE112855) (**Fig 1B, S1F-J**). Further, we examined the EMT-ness of individual CTCs and clusters of CTCs isolated from xenograft models as well as breast cancer patients [31] (GSE111065). Interestingly, while the EMT-ness of individual CTCs varied more along a spectrum, the EMT scores of CTC clusters followed a more bimodal distribution with a large difference in corresponding EMT-ness (**Fig 1C-D, S2D-I, S3A-J**). Put together, these results suggested that CTCs either freshly isolated from patients or established in culture as cell lines showed considerable heterogeneity in their EMT scores as assessed via these three independent EMT metrics. Interestingly, the EMT status of CTCs was also found to be different depending on cultured in petridish (2D) vs. in a 4D ex vivo model for a lung cancer cell line [32] (GSE50991) (**Fig 1G**).

**Figure 1.**
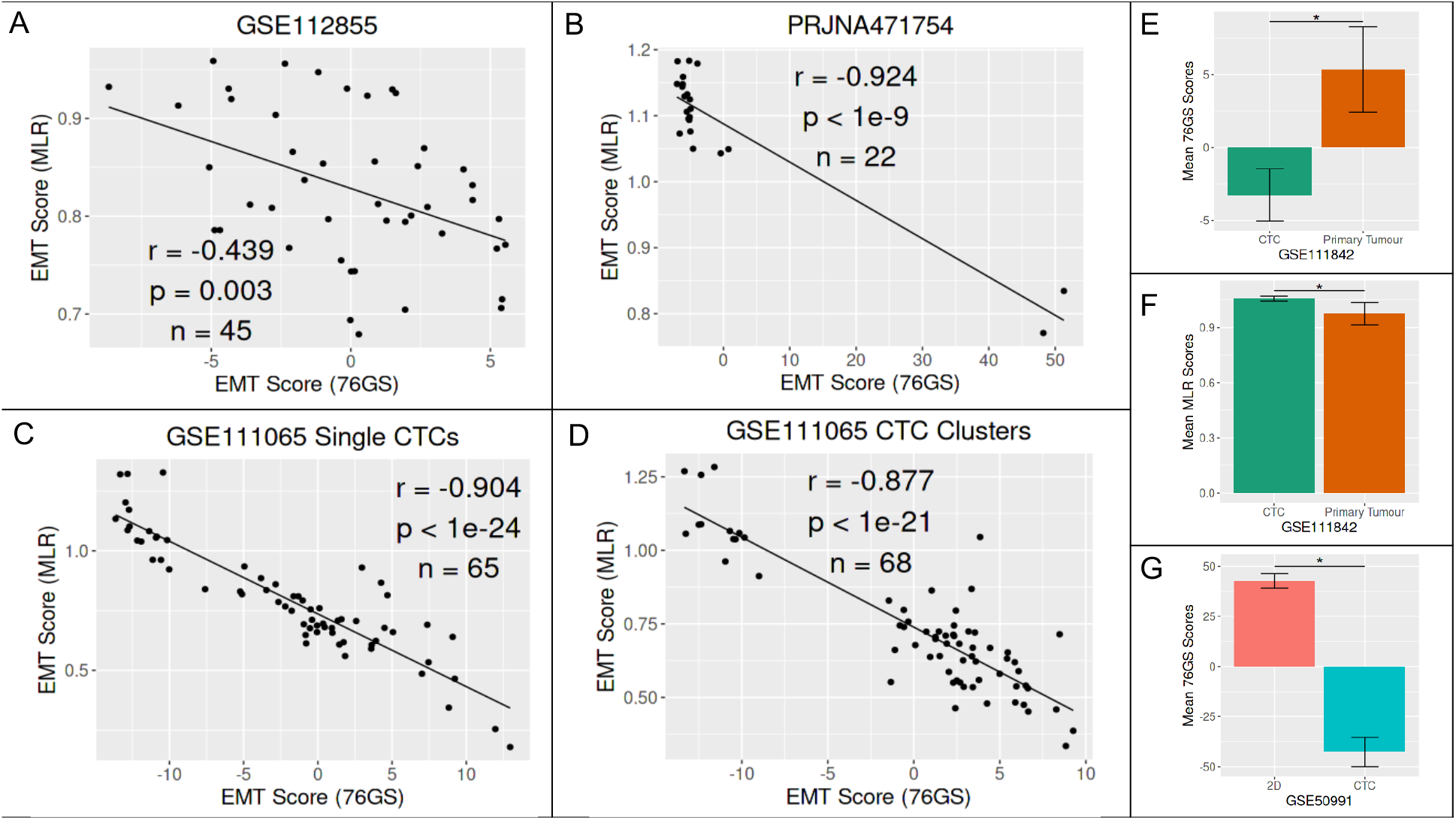
Heterogeneity in EMT scores of CTCs and primary tumors, calculated from corresponding transcriptomic data (publicly available microarray or RNA data). **(A)** Correlation between 76GS and MLR scores for GSE112855 (CTC-derived cell lines/sub lines in breast cancer) dataset. Each dot shows a sample in the dataset. Pearson’s correlation values (r, p) are reported. n = number of samples. **(B)** Same as A) but for individual CTCs in PRJNA471754. **(C**,**D)** Same as A) but for single CTCs and CTC clusters separately in GSE111065. **(E**,**F)** Comparison of EMT scores (KS, MLR) for CTC and primary tumor for GSE111842 (*: p < 0.0001). **(G)** Comparison of 76GS EMT scores for GSE50991 for 2D culture vs. 4D culture of CTCs isolated from A549 lung cancer cells (*: p < 0.0001). Students’ two-tailed t-test was used in panels E, F, G.

We also calculated the EMT scores of CTCs isolated from 16 patients in Stage II-III breast cancer and primary tumor available from 12 of them [33], and found CTCs to be relatively more mesenchymal than primary tumors (GSE111842) (**Fig 1E-F**). This observation prompted us to interrogate if the EMT status of primary tumor and CTCs can be informative of one another, i.e. can one predict the EMT status of primary tumor based on that of CTCs or *vice-versa*?

### 2. A biophysical model to investigate EMT heterogeneity

Transcriptomic analysis of tumor samples and CTCs highlighted heterogeneity in EMT scoring. Therefore, we further sought to understand the relationship between EMT-ness of primary tumor and CTCs using a simple coarse-grained biophysical model that couples phenotypic transitions driven by EMT with cell migration in the primary tumor [34].

In this model, cells are arranged on a lattice that represents the invading edge of a tumor. Based on the heterogeneity in EMT scores observed in CTCs, we considered a model with three intermediate states, E-like, E/M, M-like that are progressively less epithelial and more mesenchymal, in addition to the ‘pure’ epithelial and mesenchymal ones (**Fig. 2A, top**).

**Figure 2.**
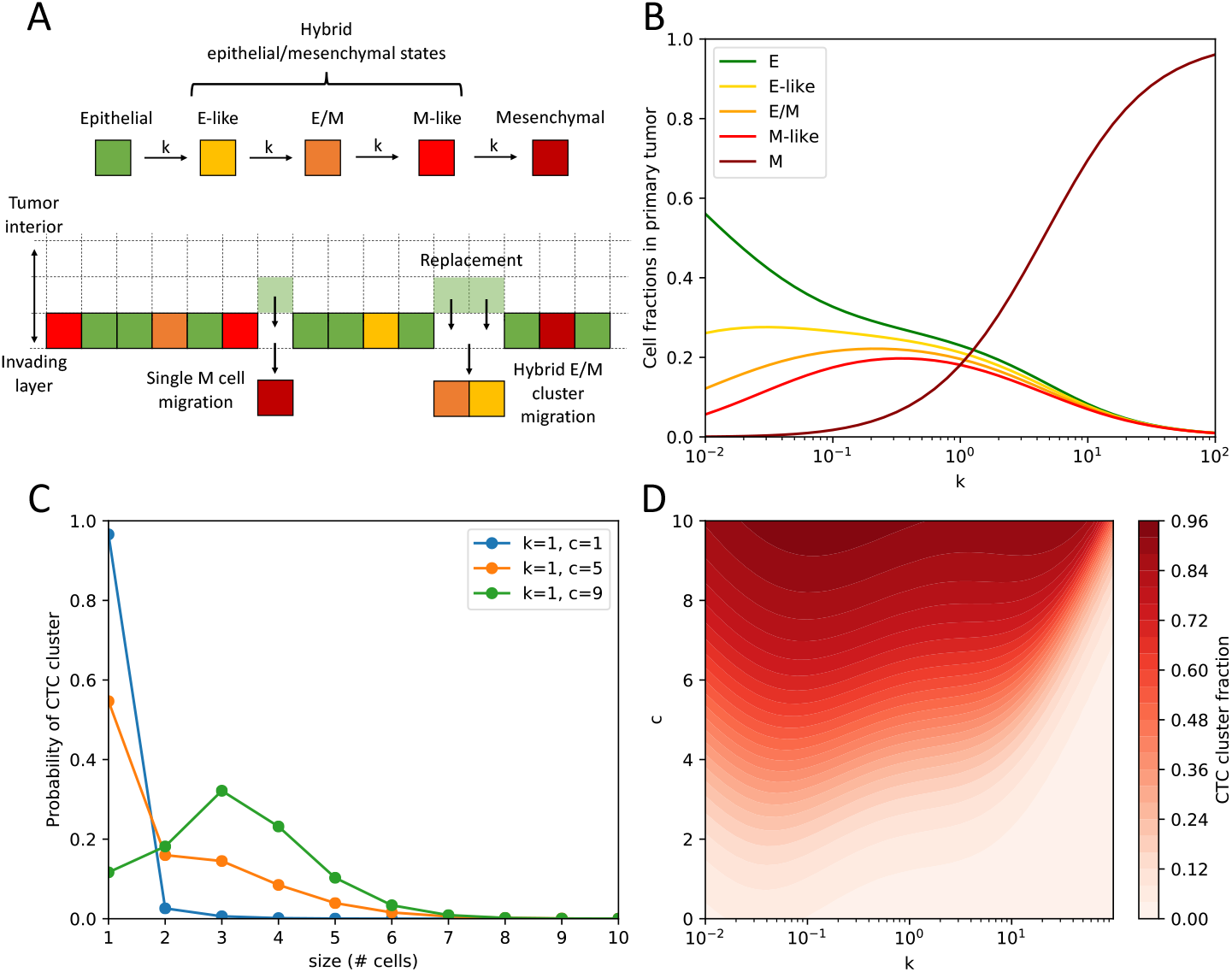
Single cell-cluster migration transition in the EMT-migration model. **(A)** Overview of the EMT-migration model. Top: cells undergo transitions through 3 hybrid E/M states with rate k. Bottom: mesenchymal cells migrate as single cells, while hybrid E/M cells can migrate together as multicellular clusters. Migrating cells are replaced by new Epithelial cells. **(B)** Fraction of Epithelial, E-like, E/M, M-like and Mesenchymal cells in the primary tumor as a function of the EMT rate (k) for a fixed value of c=2.5. **(C)** Three predicted CTC cluster size distributions for a fixed EMT rate (k=1) and increasing migration cooperativity (c). **(D)** Fraction of CTC clusters as a function of EMT rate (k) and migration cooperativity (c).

Moreover, cells undergoing EMT can migrate from the lattice. While mesenchymal cells are assumed to migrate only as single cells, hybrid E/M cells in the E-like, E/M, and M-like states can migrate together as multicellular clusters if in spatial proximity (**Fig. 2A, bottom**). The output of the model is the steady state fractions of cells in various EMT phenotypes in the tumor (E, E-like, E/M, M-like, M) as a function of the two main model’s parameters: the EMT rate (k), which describes the speed of EMT transitions, and migration cooperativity (c), which describes the propensity of hybrid E/M cells to migrate together as multicellular clusters.

The relaxation to steady state depends on two timescales: the EMT rate (k) and the migration rate, which is fixed to unit value and never varied in the model (see Methods – EMT model). Starting from an initially epithelial lattice, the model relaxes to its steady state with a speed that depends on the parameter choice (**Fig S4**). Considering that the timescales for EMT and cell migration are fast compared to the typical timescales of tumor progression [35], from now on we will always analyze the model at steady state.

We observed that varying the rate of EMT (k) at fixed cooperativity (c) can change the steady state fraction of cells with distinct EMT phenotypes in the tumor lattice: while cells are mostly epithelial for low values of k, more and more cells convert to the E-like, E/M, and M-like states as the value of k increases. Eventually, most cells are mesenchymal for large k (**Fig. 2B**). Moreover, fixing k and increasing c leads to a change in the migration strategy. At low c, the size distribution of escaping CTCs is dominated by single cells; conversely, larger values of c lead to an increased probability to observe multicellular clusters of 2 or more cells (**Fig. 2C**). More generally, varying both parameters triggers a transition from single cell migration to clustered cell migration, as seen for instance by inspecting the fraction of escape event of clusters of 2 or more cells, or cluster escape fraction (**Fig. 2D**), as well as changes in the steady state cell fractions (**Fig S5**).

Next, we investigated the implications of this model in the context of tumor-CTC EMT score relationship to provide a rationale for the observed EMT score heterogeneity in tumor and CTCs.

### 3. Collective cell migration can lead to non-trivial tumor-CTC score relationship

To investigate the relation between EMT progression in the primary tumor and CTCs, we define an EMT scoring metric in the biophysical model that can be directly compared to the scores computed in cancer datasets. Specifically, we define a metric (*S*) that can be directly compared to the MLR score. This choice is motivated by the observation that unlike the 76GS and KS metrics [13,14], the MLR score specifically focused on identifying a hybrid E/M signature [15]. Mimicking the MLR score, cells on the EMT spectrum are assigned weights ranging from 0 (epithelial) to 2 (mesenchymal). Thus, for a 5-state model, the weights for E, E-like, E/M, M-like, M states are (*w*_*E*_, *w*_*H* 1_, *w*_*H* 2_, *w*_*H* 3_, *w*_*M*_) = (0, 0.5, 1, 1.5, 2). The tumor EMT score is defined as a weighted sum of the steady state fractions of cells: *S*_*T*_ = *w*_*E*_*ρ*_*E*_ + *w*_*H*1_*ρ*_*H*1_ + *w*_*h*2_*ρ*_*H*2_ + *w*_*H*2_*ρ*_*H*3_ + *w*_*M*_*ρ*_*M*_. Similarly, a score can be defined for the CTCs by considering the fractions of migrating cells with different EMT phenotypes (*φ*_*H*1_, *φ*_*H*2_, *φ*_*H*3_, *φ*_*M*_): *S*_*CTC*_ = *w*_*H*1_*φ*_*H*1_ + *w*_*h*2_*φ*_*H*2_ + *w*_*H*3_*φ*_*H*3_ + *w*_*M*_*φ*_*M*_. Notably, since E cells cannot migrate in our model formulation, *S*_*CTC*_only considers three hybrid states and the mesenchymal state.

The tumor score (*S*_*T*_) is very close to zero (i.e. strongly epithelial) for models with low EMT rate (*k* ≪ 1, **Fig 3A**, left), and continuously increases to hybrid E/M and mesenchymal for increasing k (**Fig 3A**, left to right). Interestingly, a larger migration cooperativity (c) increases the propensity of hybrid E/M cells to undergo cluster-based migration before transitioning to a mesenchymal state, thus decreasing the tumor score (**Fig 3A**, bottom to top). Similarly, the CTC score (*S*_*CTC*_) increases with k because more cells undergo a complete EMT before migrating, and decreases with c because cells tend to migrate collectively as hybrid E/M rather than as single mesenchymal cells (**Fig 3B**). For most (k, c) parameter combinations, CTCs have a larger EMT score than the tumor (**Fig 3C**, red-shaded region). Strikingly, a condition of fast EMT and high migration cooperativity (i.e. large k, c) gives rise to a switch where *S*_*CTC*_ < *S*_*T*_ (**Fig 3C**, blue-shaded region). Overall, the relationship between *S*_*T*_ and *S*_*CTC*_ is not a fixed one but depends on (k, c), i.e. more heterogeneous, thus making it difficult to predict one from the another (**Fig 3D**). Generally speaking, CTCs are more mesenchymal than primary tumor in models with low EMT rate; conversely, CTCs can either be more or less mesenchymal than primary tumor at high EMT rate depending on the value of the migration cooperativity (c). This dependence on the model’s parameters can be observed in models with variable number of intermediate states, indicating that it represents a robust feature (**Fig S6**). From a clinical standpoint, this observation underscores the difficulty to fully characterize an invading tumor in terms of EMT when only considering samples from primary tumor or vice-versa. These results suggest that different cancer types, subtypes, and potentially even patients, might lie in different regions of the score diagram shown below.

**Figure 3.**
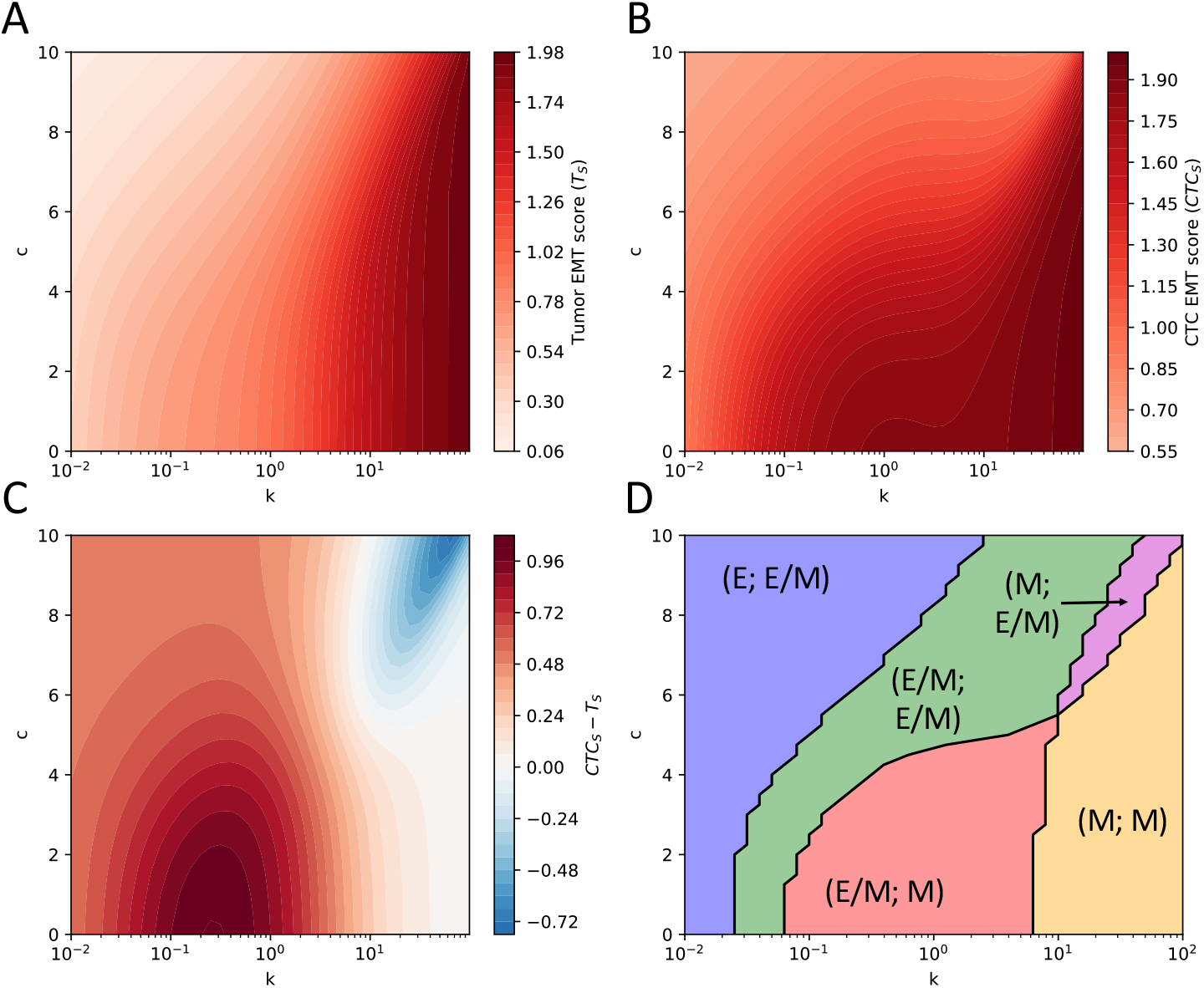
Inversions in tumor and CTC EMT scoring metrics. Predicted EMT score of primary tumor *T*_*S*_ **(A)**, migrating cells, defined as both single CTCs and CTC clusters *CTC*. **(B)**, and difference between them *CTC*_*S*_ − *T*_*S*_ **(C)** as a function of EMT rate (k) and migration cooperativity (c). **(D**) Possible combinations of EMT scores for tumor and CTCs. The scores are classified as Epithelial (*s* < 0.5), hybrid E/M (0.5 < *s* < 1.5) or mesenchymal (*s* > 1.5). For instance, (E, E/M) indicates that the tumor has an Epithelial score and the CTCs have a hybrid E/M score, and so forth.

### 4. Analysis of CTC cluster size distribution reveals variability of tumor and CTC scores across cancer types

Our EMT biophysical model predicts a heterogeneous relationship between the EMT score of primary tumors and that of CTCs. To investigate this prediction, we analyze several CTC cluster size distributions obtained experimentally through the lens of our model. In these datasets, which were obtained from different cancer types, single CTCs and CTC clusters were isolated with various techniques to obtain a frequency count or probability to observe CTC clusters with variable number of cells [36–42]. Specifically, we consider eight separate datasets isolated from either mouse models or patients from melanoma, glioblastoma, myeloma, ovarian, prostate and breast cancer [36–42]. By fitting the model’s CTC size distribution to the experimental distributions, we identify the parameter combinations (k, c) that can best fit corresponding experimental data (**SI section 1.6**). Plotting the position of the datasets on the (k, c) plane against the model’s score diagram highlights a striking heterogeneity in terms of positioning of these datasets (**Fig 4A**). In three datasets measured from melanoma, ovarian and prostate cancer, the CTCs are predicted to be considerably more mesenchymal than their corresponding tumor. However, in three other datasets from breast cancer and glioblastoma, this difference is less pronounced. Finally, two datasets from breast cancer and myeloma, respectively, are predicted to fall into the ‘inversion’ region where CTCs are less mesenchymal than the tumor (**Figure 4B**). Therefore, the model predicts that the association between EMT-ness in a primary tumor and CTCs can depend on the tumor type. Intriguingly, even within same tumor, there seems to be no generic trend in terms of EMT scores of CTCs and primary tumors, as seen in the three datasets all from breast cancer models.

**Figure 4.**
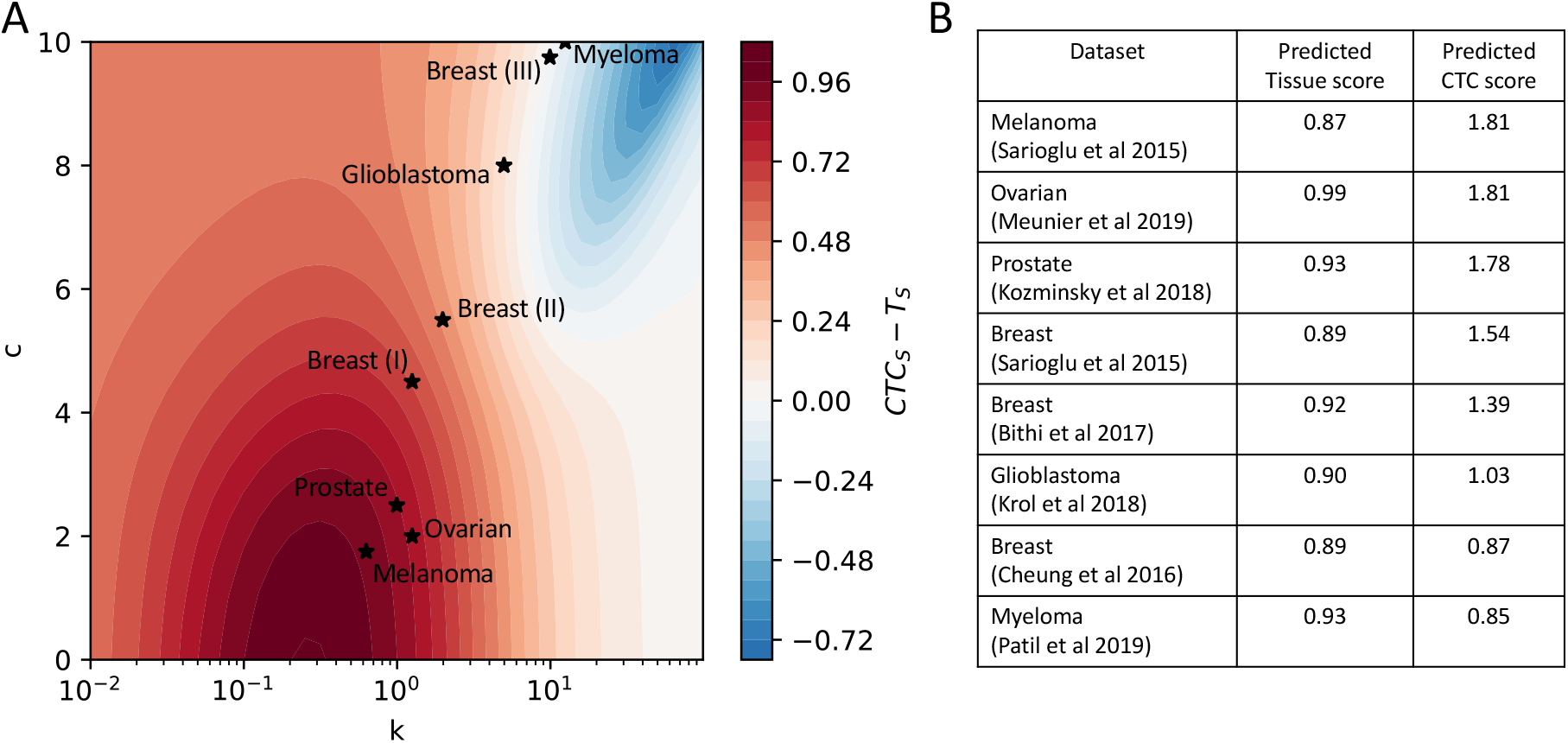
Heterogeneous tumor-CTC EMT score relationships in several CTC size distributions across cancer types. **(A)** best model fit on the (k,c) parameter space for various CTC cluster size distributions [36–42]. Each starred dot represents a CTC cluster size distribution that was fitted with the 5-state model; the x- and y-coordinates indicate the (k,c) parameter combination yielding the best fit defined in terms of minimal root square distance between experimental distribution and model prediction. **(B)** predicted tumor and CTC scores based on model’s fit.

Adding to this complexity, some of these CTC cluster size distributions consider patients with different treatment regimes. For instance, an ovarian cancer dataset (Meunier and collaborators [37]) includes patients pre- and post-chemotherapy; a prostate cancer dataset (Kozminsky and collaborators [38]) contains data from patients exposed to several different hormone therapies; and a glioblastoma dataset includes patients treated with a microtubule inhibitor (Krol and collaborators [40]). Therefore, responses to different drugs could potentially represent an additional axis of variability in the EMT score relationship between tumors and CTCs.

### 5. Exploring the predictive power of several measurements that quantify tumor aggressiveness

Motivated by the non-trivial relation between EMT scores of tumor and CTCs predicted by the model, we reviewed various types of measurements typically used to estimate tumor aggressiveness. These include EMT scores of tumors and CTCs that can be computed from single cell gene expression measurements, CTC cluster fraction in circulation and full CTC cluster size distribution. We constraint the model with each of these assays to investigate how many of the other variables can be predicted using our model (**Fig 5A**).

**Figure 5.**
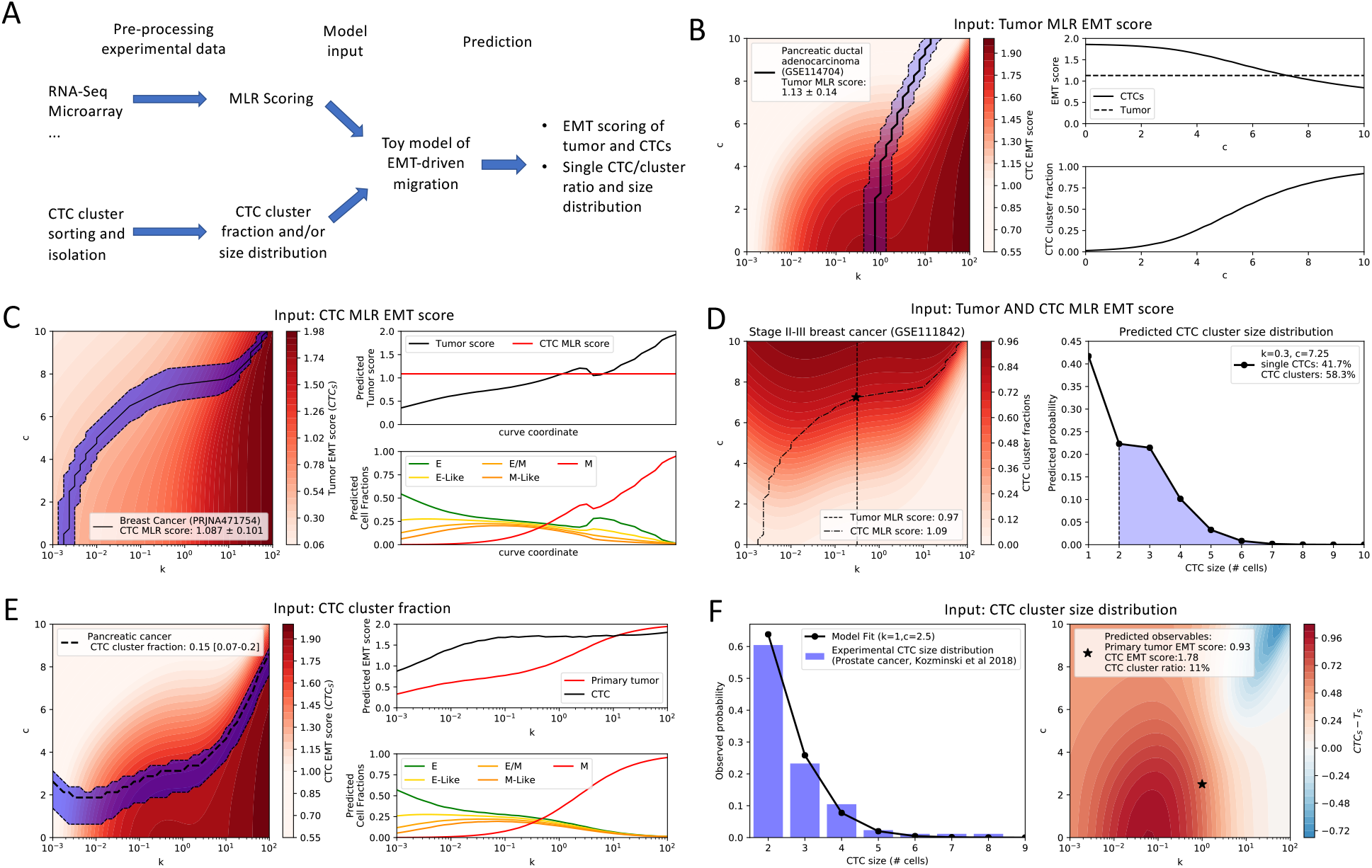
The model’s predictive power for several popular measurements of cancer aggressiveness. **(A)** For single cell measurements of gene expression (RNA-seq, microarray, etc…), an EMT scoring technique such as MLR is used to provide an input to the model; conversely, for the case of CTC cluster isolation, the input is the CTC fraction or size distribution. **(B)** left: the black line shows the contour line where the tumor EMT score is fixed and equal to the dataset’s score (the blue shading shows the standard deviation of the measurement). Right: EMT scores (top) and CTC cluster fractions (bottom) moving along the contour line. **(C)** left: the black line shows the contour line where the CTC EMT score is fixed and equal to the dataset’s score (the blue shading shows the standard deviation of the measurement). Right: EMT scores (top) and cell fractions (bottom) moving along the contour line. **(D)** left: the black lines show the contour line where the tumor and CTC EMT scores are fixed and equal to the dataset’s scores. Right: predicted CTC cluster size distribution and cluster fraction for the parameters identified in the left panel. **(E)** Left: the black line represents a model’s contour line where the CTC cluster fraction equals the experimental measurement (the blue shading shows the extremal levels found in the experiment). Right: models along the contour line exhibit variable EMT scores of tumor and CTCs (top) as well as variable cell phenotype fractions (bottom). **(F)** Left: fitting a CTC cluster size distribution provides the optimal (k,c) parameter combination. Right: predicted EMT scores and CTC cluster fraction from the model with optimal (k,c).

First, we consider the case where the EMT score of the tumor is known; this is a typical scenario if gene expression from a tumor sample is analyzed, either at a single-cell or bulk resolution. For instance, we calculate the MLR scores of several samples from a pancreatic ductal patient-derived xenograpt model [43]; on average, the samples are predicted to be hybrid E/M (1.13±0.14). In the model, constraining the tumor EMT score to a fixed value is equivalent to selecting a contour line on the two-dimensional (k, c) parameter space (**Fig 5B**, left). Each parameter combination along the curve corresponds to a model where the tumor EMT score equals the EMT score for the given dataset. Models along the contour line exhibit variable CTC EMT score and CTC cluster fraction, as seen by overlapping the contour line onto the CTC score diagram (**Fig 5B**, left). Thus, information about tumor EMT score only is not sufficient to predict neither the EMT scores of CTCs nor cluster size distribution of CTCs (**Fig 5B**, right).

Similarly, the CTC EMT score is not sufficient to fully constrain the model’s parameters. For example, we find that the average MLR score of CTCs from a breast cancer dataset [30] falls within the hybrid E/M range (1.087±0.101). The contour line at constant CTC EMT score, however, crosses parameter regions with variable tumor EMT scores (**Fig 5C**, left). Therefore, for the given CTC EMT scores, it is possible that the tumor has either a lower score (i.e. more epithelial) or higher score (i.e. more mesenchymal) that the CTCs; moreover, it can have variable cell fractions too (**Fig 5C**, right). Moreover, three other CTC datasets from prostate, myeloma and breast cancer [44–46] are mapped onto distinct model contour lines (**Fig S7A**).

Interestingly, measuring both tumor and CTC MLR score allows to identify a single (k, c) parameter combination. To illustrate this scenario, we consider a cohort of stage II-III breast cancer patients comprising RNA-sequencing of both primary tumor and CTCs [33]. The MLR metric predicts hybrid E/M signature for both tumor (0.97) and CTCs (1.09). In the model, fixing both scores corresponds to identifying two contour lines (**Fig 5D**, left); their intersection provides the (k, c) parameter combination that better reproduces the dataset. With a unique (k, c) combination, the model is able to predict a CTC cluster size distribution characterized by almost 60% of CTC clusters with 2 or more cells (**Fig 5D**, right). Unfortunately, the lack of information on the CTC cluster size distribution for this dataset prevents a comparison between model and experiment.

Another popular measurement is the overall fraction of CTC clusters, which can be obtained from blood samples by separating single CTCs and CTC clusters. Similar to the cases of tumor and CTC scores, however, a line of constant CTC fraction can be identified on the (k, c) parameter space (**Fig 5E**, left). For instance, CTC clusters isolated from a cohort of pancreatic cancer patients amount on average to 15% of the total CTCs [47]. Models along this CTC cluster fraction contour line, however, can exhibit variable tumor scores in the E, E/M and M ranges, as well as E/M or M CTC cluster scores (**Fig 5E**, right). Similarly, 4 very different CTC cluster fraction measurements are mapped onto different contour lines (**Fig S7B**) [47–51].

Finally, the model’s best fit for a complete CTC cluster size distribution can identify a unique parameter combination (k, c), thus predicting the EMT score of tumor and CTCs. In this example, a distribution from prostate cancer [38] is predicted to have a hybrid E/M tumor EMT score but a mesenchymal CTC EMT score (**Fig 5F**). Similar to the case of Fig. 5D, where the CTC cluster distribution was predicted based on EMT scores, the lack of a complete dataset with knowledge of both EMT scores and cluster size distribution prevents a quantitative model validation.

Overall, our model is capable of characterizing a tumor in terms of its EMT-ness and the propensity to undergo collective cell migration when we are aware of either both the EMT score of tumor and CTCs or the full CTC cluster size distribution. Certainly, future experimental investigations capable to estimate all the three abovementioned observables will offer a chance to test the model’s prediction more quantitatively.

## Discussion

Phenotypic heterogeneity in EMT is emerging as a hallmark of metastatic progression, and decoding its dynamics in a quantitative and predictive manner can lead to fundamental insights about metastasis [52]. Subsets of cells with varying EMT-ness (epithelial, hybrid E/M and mesenchymal states) have been observed in primary tumors, CTCs, and metastases [9,11,27].

To quantify EMT phenotypic heterogeneity, we first analyzed data from multiple tumor and CTC samples with three different EMT scoring metrics that rely on different gene lists and algorithms [13–15]. Tumors and CTCs exhibited a strong EMT score heterogeneity, suggesting that there may not exist a specific region of the EMT spectrum that is uniquely associated with tumor progression and CTC migration.

Motivated by the variability across the EMT spectrum registered in tumor tissues and CTCs, we integrated the transcriptomic analysis with a mechanism-based biophysical model that couples EMT with cell migration [34]. This model predicts a heterogeneous relation between EMT status of primary tumor and CTCs; CTCs are more mesenchymal than the primary tumor in parameter regions where invasion is mostly carried by individual cells; a transition to multicellular, clustered cell migration, however, can give rise to CTCs equally or even less mesenchymal than their corresponding tumor. In other words, measuring the level of EMT progression in CTCs does not directly allow to predict the EMT phenotypic distribution of the corresponding tumor, and *vice versa*. Specifically, in models with low EMT rate, CTCs are always more mesenchymal than primary tumor; conversely, in models with high EMT rate, CTCs can be either more or less mesenchymal than primary tumor based on the solitary or collective migration strategy, respectively. We have previously showed that high EMT rates describe well the EMT phenotype distribution of pre-treatment breast cancer patients, whereas lower EMT rates better describe the same patients after successful treatment [11,34]. Therefore, it is possible that the relation between EMT score of tumor and CTCs is not just specific to tumor type, but also evolves during cancer progression. This interesting prediction could be further explored with data from both tumor and CTCs at different stages of clinical treatment.

Nonetheless, we acknowledge multiple directions where the current model can be further improved. First, EMT transitions and migration are described with phenomenological parameters that are not explicitly connected with signaling and biophysical cellular processes. Developing models that more explicitly integrate the signaling and mechanical aspects of EMT and cell migration is a crucial future challenge, which would make the modeling even more predictive [53]. Moreover, several assumptions were made about the dynamics of cancer cell migration in order to decrease the complexity of the model. First, it is assumed that only cells at the periphery of the tumor can undergo cell migration, even though it is possible for interior cells to intravasate [54,55]. A more detailed representation of the tumor structure could more correctly account for EMT heterogeneity not only at the tumor’s invading edge, but also in more interion regions. Moreover, additional effects that could modulate migration and intravasation, such as cell death and/or breakup of multicellular clusters are not explicitly considered in the model [56]. Finally, epithelial-mesenchymal plasticity is not necessarily regulated in a cell-autonomous manner, but rather depends on communication with other cancer cells via contact-dependent signaling, such as Notch, or paracrine signaling, such as TGF-*β* [57,58]. In silico modeling of EMT and Notch underlying circuitry dynamics recently predicted that lateral induction between hybrid E/M cells can facilitate the formation of hybrid E/M multicellular clusters [59,60]. Similarly, reconstruction of TGF-*β* cell-cell communication networks recently suggested that hybrid E/M phenotypes can act as both senders and receivers of the signaling, thus facilitating EMT transitions in other cells [8]. Therefore, modeling the effect of cell-cell communication on EM plasticity and CTC cluster distributions could be another interesting future direction.

EMT is connected to multiple axes of cancer progression, including invasion, stemness and immune response [61]. How the heterogeneity along the EMT axis propagates, and in turn depends, on other hallmarks of cancer progression, remains largely unknown. Future investigations through high-throughput single cell techniques and computational modeling will help answer these questions and identify the defining principles of dynamics of EMP.

## Conclusions

As our ability to perform quantitative measurements at the level of cell migration and single-cell gene expression during metastasis increases, we need to integrate these biochemical and biophysical aspects to decipher the hallmarks of metastasis-initiating cells. While these techniques enable us to inspect EMP at unprecedented resolution, how the EMP traits of tumors propagates to those of migrating CTCs remains poorly understood. Overall, our integrated transcriptomic and computational pipeline highlights that the relationship between ‘EMT-ness’ in primary tumors and CTCs is likely heterogeneous, underscoring the need for a broader and multi-faceted approach to characterize tumor aggressiveness in the clinic.

## Acknowledgement

We thank Prof. Josè Onuchic (Rice University) for sharing his useful suggestions about the manuscript.

## Author contribution

FB and MKJ designed the research; SM and TT performed EMT score calculations; FB developed *in silico* modeling; FB and MKJ wrote the manuscript; MKJ supervised the research.

