## Supplementary information and figures for "Investigating epithelial-mesenchymal heterogeneity of tumors and circulating tumor cells with transcriptomic analysis and biophysical modeling"

**Federico Bocci[1,#,a], Susmita Mandal[2], Tanishq Tejaswi[2,3], Mohit Kumar Jolly[2,#]**

[1]Center for Theoretical Biological Physics, Rice University, Houston, TX;
[2]Centre for BioSystems Science and Engineering, Indian Institute of Science, Bangalore, India
[3]UG Programme, Indian Institute of Science, Bangalore, India;
[a]Current address: NSF-Simons Center for Multiscale Cell Fate Research, University of California, Irvine, CA

### 1. Mathematical model of EMT and migration

Here, we provide the details of the EMT mathematical model: The model was explicitly derived for the case of one intermediate state; in the following sections, the same derivation is provided for the case of 3 intermediate states. Generalizations to a different number of intermediates can be obtained with a similar procedure.

#### 1.1 Equations for the case of three intermediate hybrid states

The temporal dynamics of the fractions of E, E-like, E/M, M-like, and M cells is described by a set of ordinary differential equations:

$$\frac{d\rho_E}{dt} = -k\,\rho_E + \Theta(\rho_{H_1},\rho_{H_2},\rho_{H_3}) + \rho_M \qquad (1a)$$

$$\frac{d\rho_{H_1}}{dt} = +k\,\rho_E - k\,\rho_{H_1} - \frac{\rho_{H_1}}{\rho_{H_1}+\rho_{H_2}+\rho_{H_3}}\Theta(\rho_{H_1},\rho_{H_2},\rho_{H_3}) \quad (1b)$$

$$\frac{d\rho_{H_2}}{dt} = +k\,\rho_{H_1} - k\,\rho_{H_2} - \frac{\rho_{H_2}}{\rho_{H_1}+\rho_{H_2}+\rho_{H_3}}\Theta(\rho_{H_1},\rho_{H_2},\rho_{H_3}) \quad (1c)$$

$$\frac{d\rho_{H_3}}{dt} = +k\,\rho_{H_2} - k\,\rho_{H_3} - \frac{\rho_{H_3}}{\rho_{H_1}+\rho_{H_2}+\rho_{H_3}}\Theta(\rho_{H_1},\rho_{H_2},\rho_{H_3}) \quad (1d)$$

$$\frac{d\rho_M}{dt} = +k\,\rho_{H_3} - \rho_M \qquad (1e)$$

In Eqs. 1, k is a dimensionless EMT rate parameter that represents the ratio between the rate of EMT and the basal rate of escape ($k = k_{EMT}/k_{ESC}$). The term $\Theta(\rho_{H_1},\rho_{H_2},\rho_{H_3})$ in eqs. (1b-d) quantifies the loss of E-like, E/M and M-like cells due to migration (either as single cells or multicellular clusters); the form of this term will be derived in the following section. In the main text, we refer to k simply as the rate of EMT for short notation. The term $-\rho_M$ in eq. (1e) represents the loss of mesenchymal cells due to single cell migration; since the migration rate of M cells is constant, this term is simply proportional to the fraction of M cells in the lattice. Finally, the lattice is replenished of new epithelial cells to substitute E-like, E/M, M-like and M cells lost due to migration; hence the term $+\Theta(\rho_{H_1},\rho_{H_2},\rho_{H_3}) + \rho_M$ in eq. (1a). The steady state cell fractions (${\rho_E}^{(eq)}$, ${\rho_{H_1}}^{(eq)}$, ${\rho_{H_2}}^{(eq)}$, ${\rho_{H_3}}^{(eq)}$, ${\rho_M}^{(eq)}$) are obtained by setting eq. (1) to zero.

#### 1.2 Escape flux of hybrid cells

We first write down the expression of the escape flux $\Theta$ in a generic setting (i.e. without assumptions on lattice geometry and dimensionality and/or specific form of adhesion terms). The escape flux of hybrid cells (E-like, E/M and M-like) per unit time considers clusters of any size composed by a mixture of E-like, E/M and M-like cells. Moreover, these multicellular clusters can break down into smaller subclusters that migrate independently. To make this approach computationally feasible, we consider a mean-field approximation of the form:

$$\Theta(\rho_{H_1},\rho_{H_2},\rho_{H_3}) = \sum_{s=1}^{+\infty} p_s(\rho_{H_1},\rho_{H_2},\rho_{H_3}) \sum_{m=1}^{s} n(m,s)\,K(m,s) \qquad (2)$$

In eq. (2), the first sum quantifies the probability to find clusters of any size s ($p_s(\rho_{H_1},\rho_{H_2},\rho_{H_3})$) given the fractions $\rho_{H_1},\rho_{H_2},\rho_{H_3}$. The second sum, at fixed cluster size (s), considers all possible subclusters of size m<s that could arise if the cluster breaks up. Therefore, the summation is a product of relative frequencies of subclusters with various sizes ($n(m,s)$) and escape rates for each one of these subclusters ($K(m,s)$).

1.3 Effective-2D approximation for Θ

The analytical expression of the cluster size distribution as a function of cell fraction cannot be established analytically in any dimension larger than two. For this reason, we consider a model where cells are arranged in an infinite one-dimensional chain. In this case, the probability to find a cluster of size s that is an arbitrary mixture of E-like, E/M, and M-like cells is $p_s(\rho_H) = {\rho_H}^s(1-\rho_H)^2$, where we defined $\rho_H = \rho_{H_1} + \rho_{H_2} + \rho_{H_3}$ for convenience.

In the one-dimensional chain, a cluster of s cells can only be a linear chain. The number of existing subclusters of size m<s within the cluster of size s is:

$$N(m,s) = s - m + 1 \quad (3)$$

The relative frequency of subclusters of a given size $n(m,s)$ is defined by normalizing the number of subclusters $N(m,s)$:

$$n(m,s) = \frac{N(m,s)}{\sum_{m=1}^{s} N(m,s)} = \frac{s-m+1}{\frac{s(s+1)}{2}} \quad (4)$$

1.4 Escape rate of hybrid clusters

The escape rate of a subcluster composed by m H-cells enclosed within a larger cluster composed by s H-cells is:

$$K(m,s) = m^c \prod_i {p_i}^{n_i} \quad (5)$$

In eq. (5), the first factor ($m^c$) describes the propensity of hybrid cells to migrate collectively; m is the number of cells n the cluster and c is a migration cooperativity index. Larger values of c imply that larger clusters have larger escape rates. The second term describes the breaking of bonds with neighboring cells that is necessary to migrate. The sum runs over all possible combinations of hybrid cells in the lattice (E-like, E/M, M-like) and neighboring cells that share an adhesive bond (E, E-like, E/M, M-like). In the simple case of one intermediate state (H), a cluster of m H cells has 2m+2 nearest neighbors. Of these, a fraction $n_E = \rho_E/(\rho_E + \rho_H + \rho_M)$ are epithelial and a fraction $n_H = \rho_H/(\rho_E + \rho_H + \rho_M)$ are H. The bonds with E neighbors are broken with probability $p_E$ and the binds with H neighbors are broken with probability $p_H$. M neighbors do not form adhesive bonds in the model, and therefore are not considered. Therefore, the product in eq. (5) amounts to ${p_E}^{n_E}{p_H}^{n_H}$. In the case of interest with 3 intermediate states, many more combinations of cells phenotypes are possible. The fractions of neighboring cells in various states at any time $t$ are given by $\rho_E(t), \rho_{H1}(t), \rho_{H2}(t), \rho_{H3}(t), \rho_M(t)$. The relative fractions of E-like, E/M, M-like cells within the cluster are:

$$h_{H1} = \frac{\rho_{H1}}{\rho_{H1} + \rho_{H2} + \rho_{H3}} \quad (6a)$$

$$h_{H2} = \frac{\rho_{H2}}{\rho_{H1} + \rho_{H2} + \rho_{H3}} \quad (6b)$$

$$h_{H3} = \frac{\rho_{H3}}{\rho_{H1} + \rho_{H2} + \rho_{H3}} \quad (6c)$$

Under the ‘effective-2D’ approximation, a subcluster of m cells has 2m+2 nearest neighbors, and thus 2m+2 pairs of cells, one belonging to the subcluster and one neighbor. The number of pairs $n_{ij}$, where $i$ is the state of the cell in the subcluster and $j$ is the state of the neighbor, is:

$$n_{ij} = (2m+2)h_i\rho_j \quad (7)$$

Where $h_i$ is the relative fraction of cells in state $i$ in the subcluster and $\rho_j$ is the fraction of cells in state $j$ in the lattice. Generalizing for all possible pairs, the product in eq. (5) becomes:

$$\prod_i p_i^{n_i} = p_{E-H1}{}^{n_{E-H1}} p_{E-H2}{}^{n_{E-H2}} p_{E-H3}{}^{n_{E-H3}} p_{H1-H1}{}^{n_{H1-H1}}\, p_{H1-H2}{}^{n_{H1-H2}} p_{H1-H3}{}^{n_{H1-H3}} \cdot p_{H2-H2}{}^{n_{H2-H2}} p_{H2-H3}{}^{n_{H2-H3}}\, p_{H3-H3}{}^{n_{H3-H3}} \quad (8)$$

Where the set of $p_{i-j}$ represent the probabilities to break adhesive bonds between cells in states $i$ and $j$. It is worth noting that, in the limit case of a single mesenchymal (M) cell, m=1 and there are no adhesion bonds, therefore eq. (5) naturally relaxes to 1.

1.5 Estimation of probabilities to break cell-cell adhesion bonds

We assume that cell states are ordered on an adhesion scale, with the E state having the maximal adhesion factor $A_E$=1 and the M state having the minimal adhesion factor $A_M$=0. The three intermediate states are placed in between in a linear fashion, so that $A_{H1}$=0.25, $A_{H2}$=0.5, $A_{H3}$=0.75 The adhesion factor between a pair of cells in states I and j, respectively, is defined as the product $A_iA_j$. $A_iA_j$ can be interpreted as the probability to maintain the bond. Therefore, the complementary $p_{ij}$=1- $A_iA_j$ is the probability to break the bond.

1.6 Fitting of experimental CTC cluster size distributions

Histograms of CTC clusters as a function of CTC cluster size were taken from published literature and normalized to unit value. The best model fit is identified by the parameter combination (k,c) yielding the minimum sum of squared distances between experimental distribution and model's prediction (all other model parameters are fixed for this calculation). For some of the experimental distributions, data was not available for single CTCs (i.e. the histogram starts at a minimum CTC size of n=2 cells). In these cases, we compared the experimental CTC size distribution with the model's distribution re-normalized to exclude single CTCs.

### Supplementary Figures

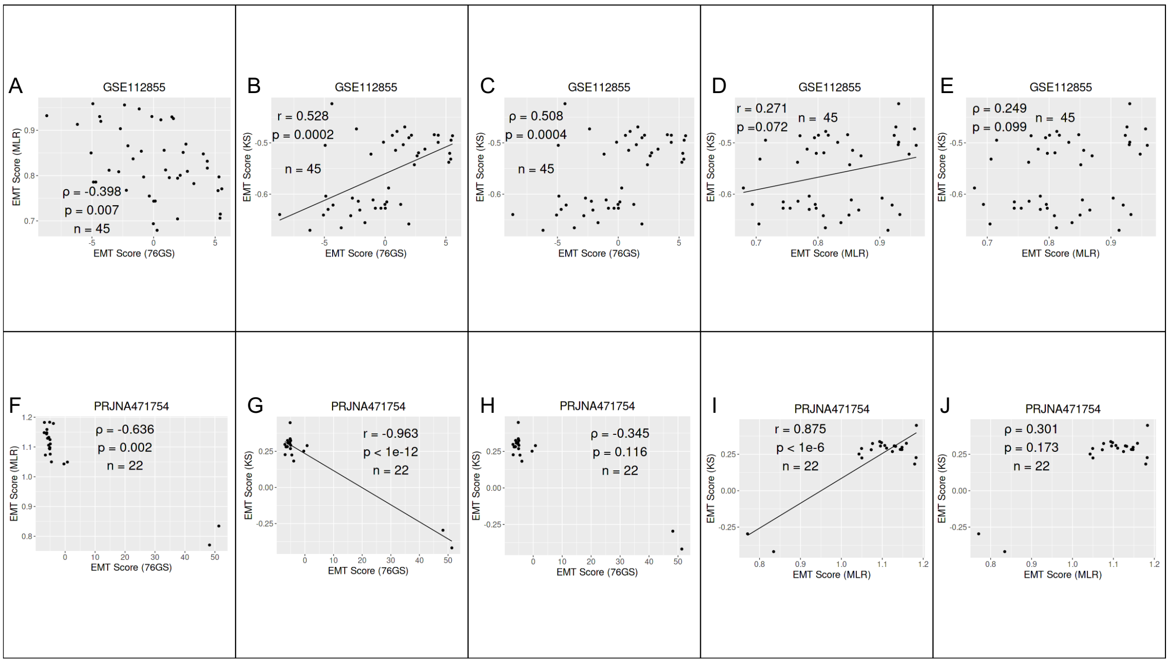


Supplementary Figure 1. Heterogeneity in EMT scores. **A)** Same as Fig 1A but showing Spearman correlation. **B, C)** Scatter plots based on Pearson and Spearman correlation for KS and 76GS scores. **D, E)** Same as B, C) but for MLR and KS scores. **F)** Same as Fig 1B but showing Spearman correlation. **G, H)** Scatter plots based on Pearson and Spearman correlation for KS and 76GS scores. **I, J)** Same as G, H) but for MLR and KS scores.

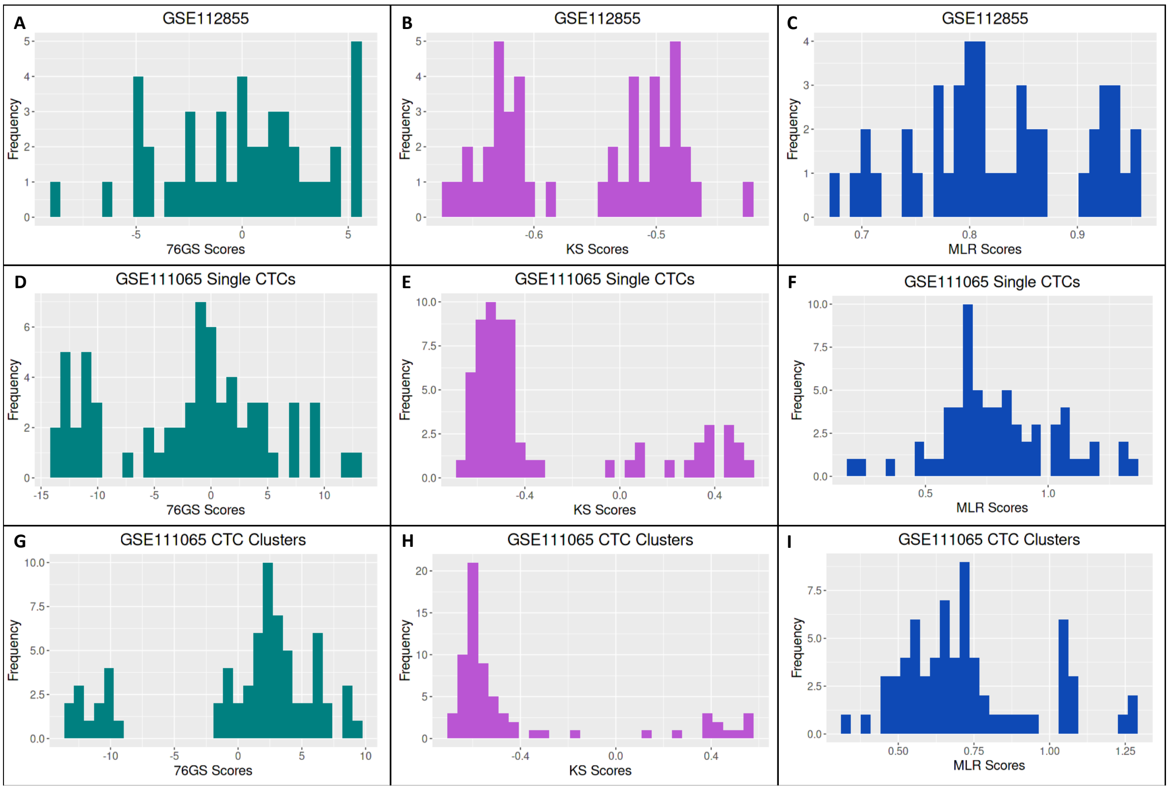


Supplementary figure 2. Histograms showing heterogeneity in EMT Scores. A) For 76GS scores of GSE112855. B) Same as A but for KS. C) Same as A but for MLR. D,E,F) Same as A-C but for GSE111065 Single CTCs. G,H,I)Same as A-C but for GSE111065 CTC Clusters.

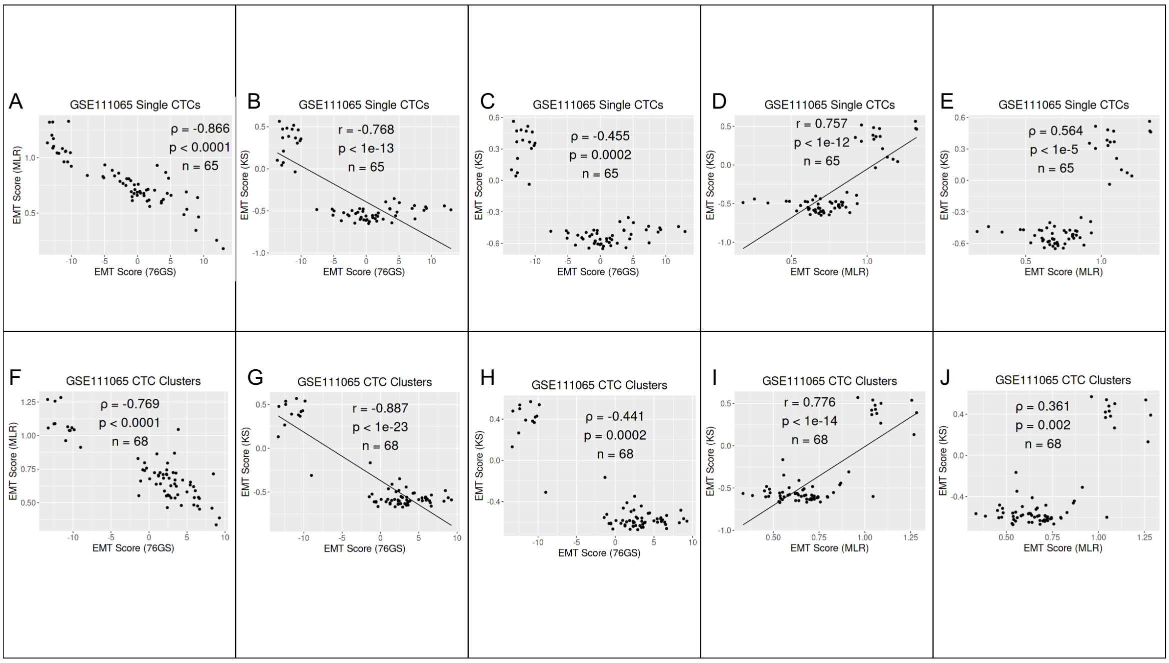


Supplementary Figure 3. Heterogeneity in EMT scores. **A)** Same as Fig 1C but showing Spearman correlation. **B, C)** Scatter plots based on Pearson and Spearman correlation for KS and 76GS scores. **D, E)** Same as B, C) but for MLR and KS scores. **F-J)** Same as panels A-E but for CTC clusters in GSE 111065 (Fig 1D) instead of single CTCs (Fig 1C).

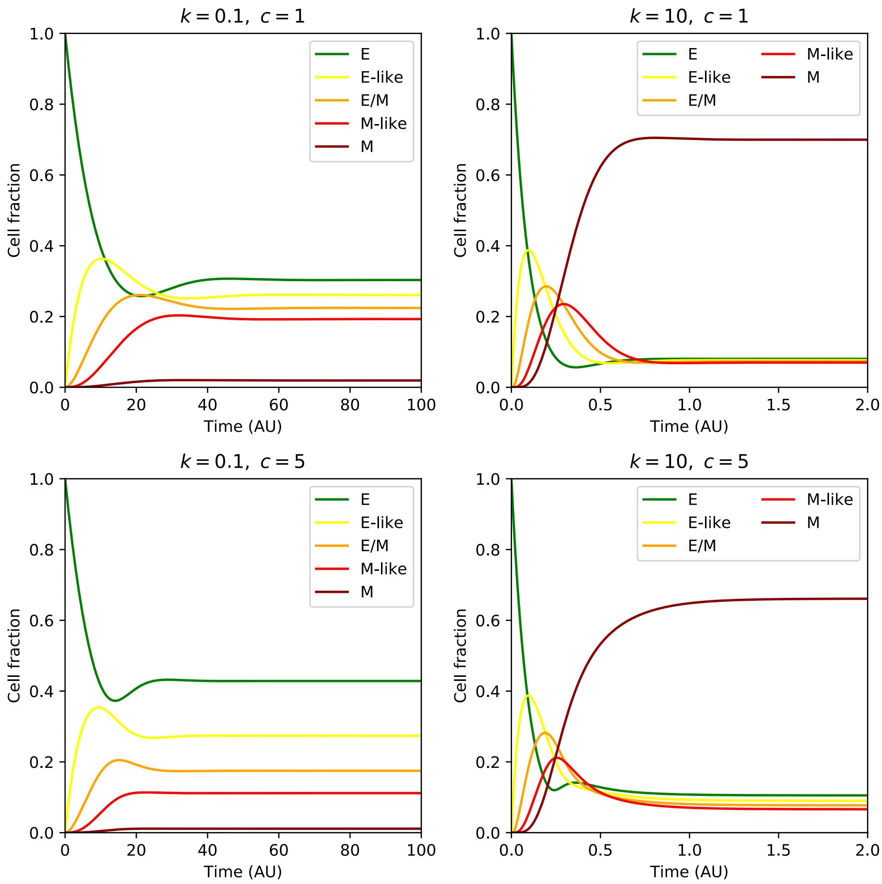


Supplementary Figure 4. Fraction of E, E-like, E/M, M-like and M cells as a function of time starting from an initial condition $\rho_E = 1$. Four panels show relaxation to steady state for four different (k,c) parameter combinations.

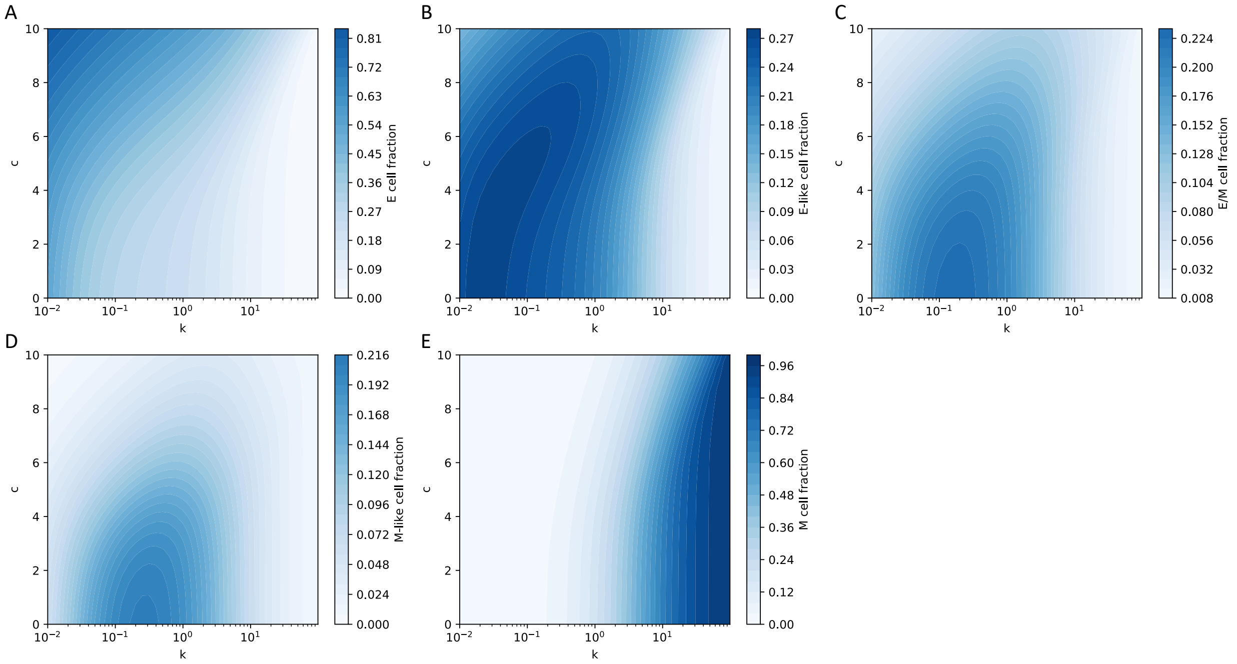


Supplementary Figure 5. Steady state fractions of E, E-like, E/M, M-like, and M cells as a function of EMT rate (k) and migration cooperativity (c).

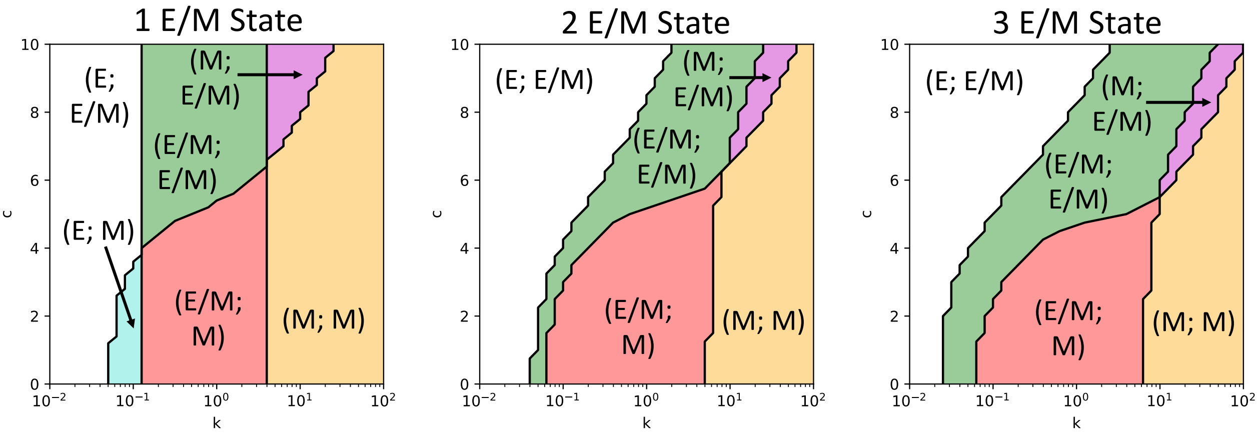


Supplementary Figure 6. Possible combinations of EMT scores for tumor and CTCs for models with one intermediate state (left), two intermediate states (center) and three intermediate states (right, same as Fig. 3D).

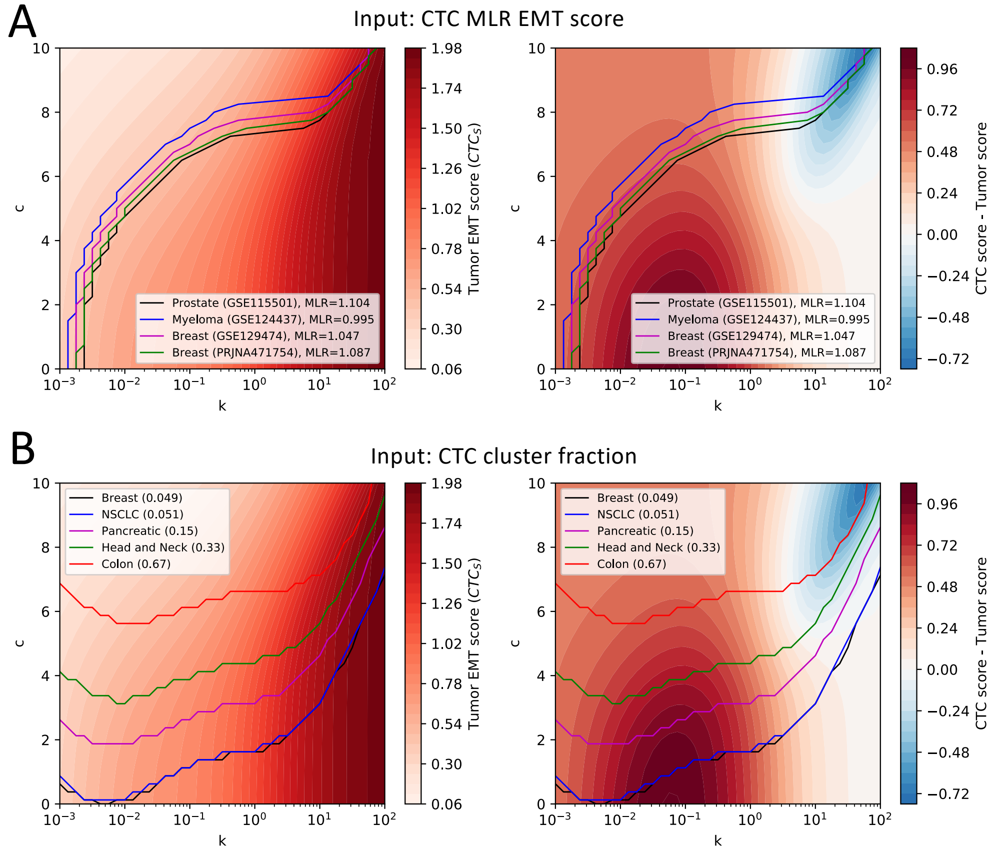


Supplementary Figure 7. **(A)** Contour lines of constant CTC EMT score for 4 independent datasets including Prostate cancer, Myeloma, and Breast cancer against (i) tumor EMT score (left) and (ii) difference between CTC score and tumor score (right). **(B)** Contour lines of constant CTC cluster fraction for 5 independent datasets including Breast cancer, Non-small cell lung cancer, Pancreatic cancer, Head and neck cancer, and Colon cancer against (i) tumor EMT score (left) and (ii) difference between CTC score and tumor score (right).
